# Protein SUMOylation is crucial for phagocytosis in *Entamoeba histolytica* trophozoites

**DOI:** 10.1101/2021.02.01.429131

**Authors:** Mitzi Díaz-Hernández, Rosario Javier Reyna, Izaid Sotto-Ortega, Guillermina García-Rivera, Maricela Sarita Montaño, Abigail Betanzos, Esther Orozco

## Abstract

During phagocytosis, a key event in the virulence of the protozoan *Entamoeba histolytica*, several molecules in concert contact the target, generate pseudopodia, and internalize and digest the ingested prey. Posttranslational modifications provide proteins the timing and signaling to intervene in these processes. SUMOylation is a posttranslational modification that in several systems grants a fine tuning for protein functions, protein interactions and cellular location, but it has not been studied in *E. histolytica*. In this paper, we characterized the *E. histolytica SUMO* gene and its product (EhSUMO) and elucidated the EhSUMO 3D-structure. Furthermore, here we studied the relevance of SUMOylation in phagocytosis, particularly in its association with EhADH (an ALIX family protein) and EhVps32 (a protein of the ESCRT-III complex), both involved in phagocytosis. Our results indicated that EhSUMO has an extended N-terminus that differentiates other SUMO from ubiquitin. It also presents the GG residues at the C-terminus and the ΨKXE/D binding motif, both involved in target protein contact. Additionally, *E. histolytica* genome possesses the enzymes belonging to the SUMOylation-deSUMOylation machineries. Confocal microscopy assays, using α−EhSUMO antibodies disclosed a remarkable membrane activity with convoluted and changing structures in trophozoites during erythrophagocytosis. SUMOylated proteins appeared in pseudopodia, phagocytic channels, and around the adhered and ingested erythrocytes. Docking analysis predicted interaction of EhSUMO with EhADH, and immunoprecipitation and immunofluorescence assays revealed that the EhADH-EhSUMO association increased during phagocytosis, whereas the EhVps32-EhSUMO interaction appeared stronger since basal conditions. In *EhSUMO* knocked down trophozoites, the bizarre membranous structures disappeared, and EhSUMO interaction with EhADH and EhVps32 diminished. Our results evidenced the presence of a *SUMO* gene in *E. histolytica* and the SUMOylation relevance during phagocytosis.

**Author’s Abstract:** Phagocytosis is one of the main functions that *Entamoeba histolyitica* trophozoites carry out during the invasion to the host. Many proteins are involved in this fascinating event, in which the plasmatic membrane undergoes to multiple and speedy changes. Posttraductional modifications activate proteins in the precise time that they must get involved. SUMOylation, that consists in the non-covalent binding of SUMO protein with target molecules, is one of the main changes suffered by proteins in order to enable them to participate in cellular functions. SUMOylation had not been studied in *E. histolytica* nor in phagocytosis, and our working hypothesis is that this event is deeply engaged in the ingestion of target molecules and cells. The results of this paper prove the presence of an intronless *bona fide EhSUMO* gene encoding for a predicted 12.6 kDa protein that is actively involved in phagocytosis. Silencing of the *EhSUMO* gene affected the rate of phagocytosis and interfered with the EhADH and EhVps32 function, two proteins involved in phagocytosis, strongly supporting the importance of SUMOylation in this event.

## Introduction

As in other eukaryotes, in *Entamoeba histolytica*, the protozoan causative of human amoebiasis, cellular activities, including the attack to the target cell, are widely controlled by posttranslational modifications (PTMs) of proteins. Together with the perpetual movement of trophozoites, virulence expression requires intensive vesicular traffic and association to and disassociation from proteins performing the concatenated events that conduct target molecules through several compartments, for recycling or digestion. PTMs range from peptide bond cleavage and addition of phosphate and other small chemical groups, carbohydrates and lipids to the alteration of proteins by the conjugation of modifiers such as ubiquitin and SUMO. The knowledge of the changes suffered by molecules involved in adherence to and invasion and phagocytosis of trophozoites to the target cells are pivotal to get a more comprehensive view of the parasite virulence mechanisms.

SUMO is a 10 to 13 kDa small ubiquitin-related modifier that shares 18% similarity with ubiquitin in its three-dimensional structure (1). In the N-terminus, SUMO has an extended 10 to 25 amino acid chain, absent in ubiquitin. As this later, SUMO conjugates to its target by an isopeptide bond formed on a lysin present in the consensus motif ΨKXE/D (Ψ: hydrophobic amino acid, K: target lysin, X: any amino acid, E and D: glutamic and aspartic acids, respectively) (2). However, many other reports indicate that alternative sequences can be used for SUMO binding (3–5). Furthermore, SUMO can associate to target proteins as a single moiety or as SUMO polymers (6) in an ATP-dependent event that requires the action of E1, E2, and E3 enzymes. DeSUMOylation, the reverse process, occurs by the specific proteases UIp1a and UIp1b (7).

SUMOylation-deSUMOylation is a switch to control the cellular location of proteins and their interaction with other molecules (8). Cell growth, differentiation, response to stress, regulation of signal transduction, gene expression, and chromatin remodeling, among others, require SUMOylation of certain proteins (9,10). In protozoan parasites, it is known that SUMOylation takes part in cell-cycle progression and influences morphology in *Giardia lamblia* (11), while in *Trypanosoma brucei*, it contributes to chromatin organization (12) and in *Plasmodium falciparum*, participates in invasion (13).

ESCRT machinery is deeply involved in the phagocytosis of *E. histolytica* trophozoites (14–16). EhVps2, EhVps20, EhVps24, and EhVps32, members of ESCRT-III complex, and EhADH, an ALIX family protein (15) and accessory member of the ESCRT machinery, participate in membrane deformation, necessary for pseudopodia and vesicle generation (17). They are also part of the scission apparatus to form intraluminal vesicles (ILVs) in multivesicular bodies (MVBs) (17). SUMOylation could be one of the signals for the time and place to the ESCRT proteins to act during ingestion and processing of the prey, but SUMO has not been characterized in *E. histolytica*. Besides the advancement on the comprehension of the molecular events occurring during phagocytosis, the importance of understanding these phenomena relies on the possibility of carrying out the blockage of specific parasite molecules and develop better diagnostic and therapeutic methods against *E. histolytica* that infects 50 million people and kills 100,000 annually, around the world (18). We pursued here the identification and characterization of SUMO in this parasite and investigated whether ESCRT proteins are SUMOylated during phagocytosis. Particularly, we explored the association of SUMO with EhADH and EhVps32 proteins to scrutinize the changes that they undergo, during this event. The results evidenced the active participation of SUMOylation in phagocytosis.

## Results

### *In silico* analysis predicts the existence of SUMO and the SUMOylation-deSUMOylation machineries in *E. histolytica*

To investigate whether the proteins involved in phagocytosis go through SUMOylation, we first performed bioinformatic analysis to search for *SUMO* genes in the AmoebaDB (http://amoebadb.org/amoeba/), using as template the *SUMO* sequence from *G. lamblia* (11,19). Our search revealed two candidates in *E. histolytica*: EHI_170060 and EHI_151620. However, EHI_151620 predicts a product without the glycine residues (GG) at the C-terminus, a characteristic of SUMO (20). In contrast, the EHI_170060 open reading frame is a 345 bp intronless sequence with 33% homology to the *G. lamblia SUMO* gene. It is located at the complementary DNA strand, between the fragments annotated as EH_170050 and EH_170070, at the 46961 and 47530 bp (Fig. 1A). By its sequence, we estimated a protein of 12.6 kDa (EhSUMO) that has the extra amino acids at the amino terminus, recognized as the principal difference between SUMO and ubiquitin (21,22). It has two ΨKXE/D consensus motifs at 21 to 24 and 30 to 34 residues, and displays the GG doublet at the C-terminus (112 and 113 residues), both described as SUMO interaction sites for target proteins (20) (Fig. 1B). *E. histolytica*, as other protozoa (11), and *Saccharomyces cerevisiae* (23), *Drosophila melanogaster* (24) and *Caenorhabditis elegans* (25), has only one intronless *SUMO* gene, while vertebrates have four (3), and plants, eight (26).

**Figure 1.**
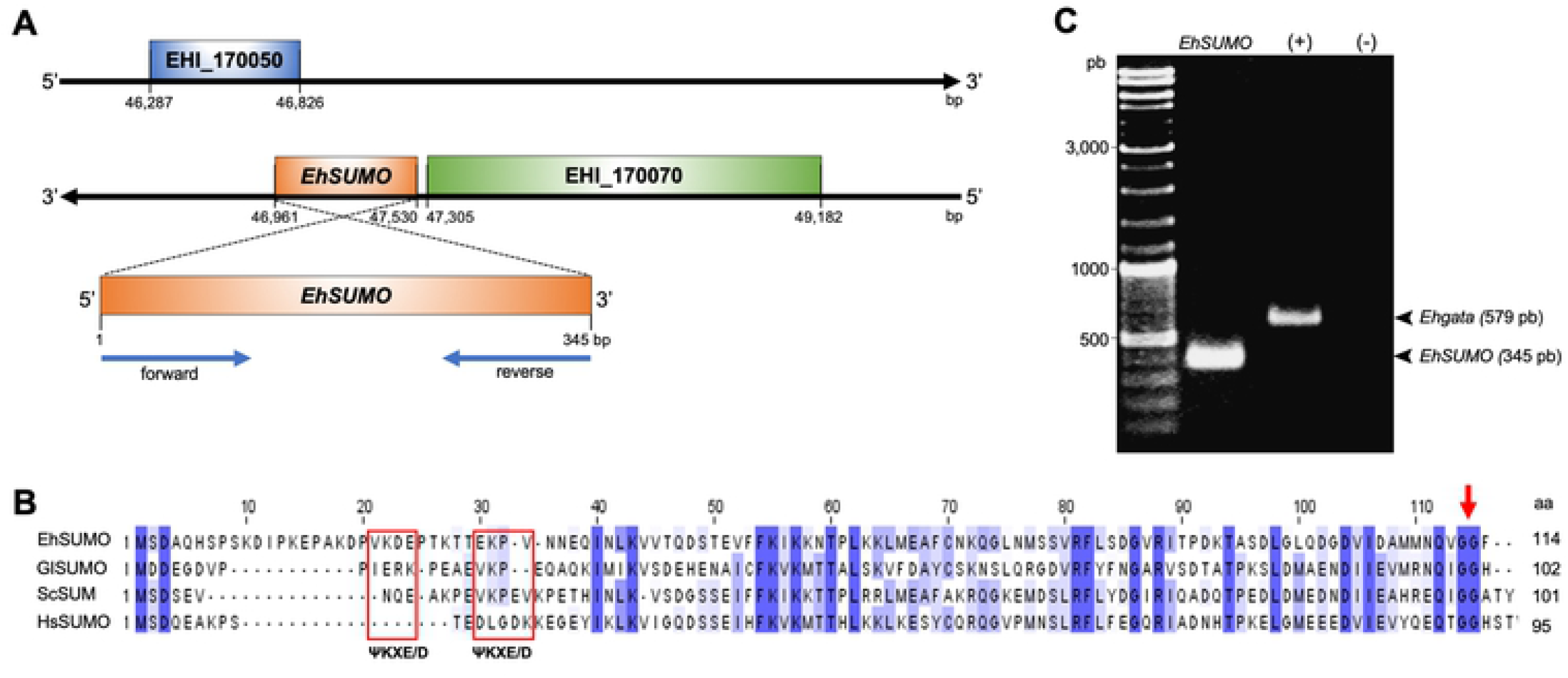
Identification and location of the *EhSUMO gene* in *E. histolytica genome*. A) Location of *EhSUMO* gene in the genome. EhSUMO in a contig located between 47,530 to 46,961 bp in the DNA negative chain flanked by two hypothetic genes (EHI_170050 and EHI_170070). Blue arrows: primer designed for *EhSUMO* amplification. B) Comparative alignment of predicted EhSUMO amino acid sequence with proteins from other organisms, the blue color indicates the identity and similarity of amino acids. Red squares signal the **ΨKXE/D** motifs and arrow the GG doublet. C) PCR amplification using complementary DNA (cDNA) as template and specific primers for *EhSUMO. Ehgata*: (+) positive control). Water (-): negative control. Arrowheads: PCR products.

Multiple alignments of EhSUMO amino acid sequence with SUMO of *S. cerevisiae, H. sapiens*, and *G. lamblia*, revealed 55, 48, and 33% identity, respectively (Table I), and the whole gene sequence confirmed the presence of the additional bases at the amino terminus and the GG motif at the C-terminus (Fig. 1C).

The interactome, carried out with the putative EhSUMO as a bait and the STRING database (http://sumosp.biocuckoo.org.), predicted *E. histolytica* proteins that interact with SUMO to perform different functions (Table II), already described in other systems. Next, we identified the putative *E. histolytica* genes and proteins required for SUMOylation, to compare them with those of other organisms (7). E1 (EHI_035540), E2a (EH1_178500), E2b (EHI_14747), E3a (EHI_0988320) and E3b (EHI_069470), presumptive SUMOylation enzymes, revealed identities from 29 to 56 % to *S. cerevisiae, 2*7.8 to 53.5 % to *H. sapiens* SUMO-2 gene, and 25 to 49% to *G. lamblia* (Table I). Putative proteins in charge of deSUMOylation: UIp1a (EHI_067510), and UIp1b (EHI_097940) exhibited 23.6 to 44.3% identities to their orthologues (Table I). The phylogenetic tree obtained using the MEGAT7 software, showed that EhSUMO is close to *D. discoideum, T. cruzi* and *Toxoplasma gondii* SUMO proteins, whereas it has a more distant phylogenetic relationship with the *H. sapiens* orthologues (Fig. 2). The bioinformatic analysis strongly suggest that *E. histolytica* has a *SUMO* gene, and those involved in the SUMOylation and deSUMOylation machineries.

**Figure 2.**
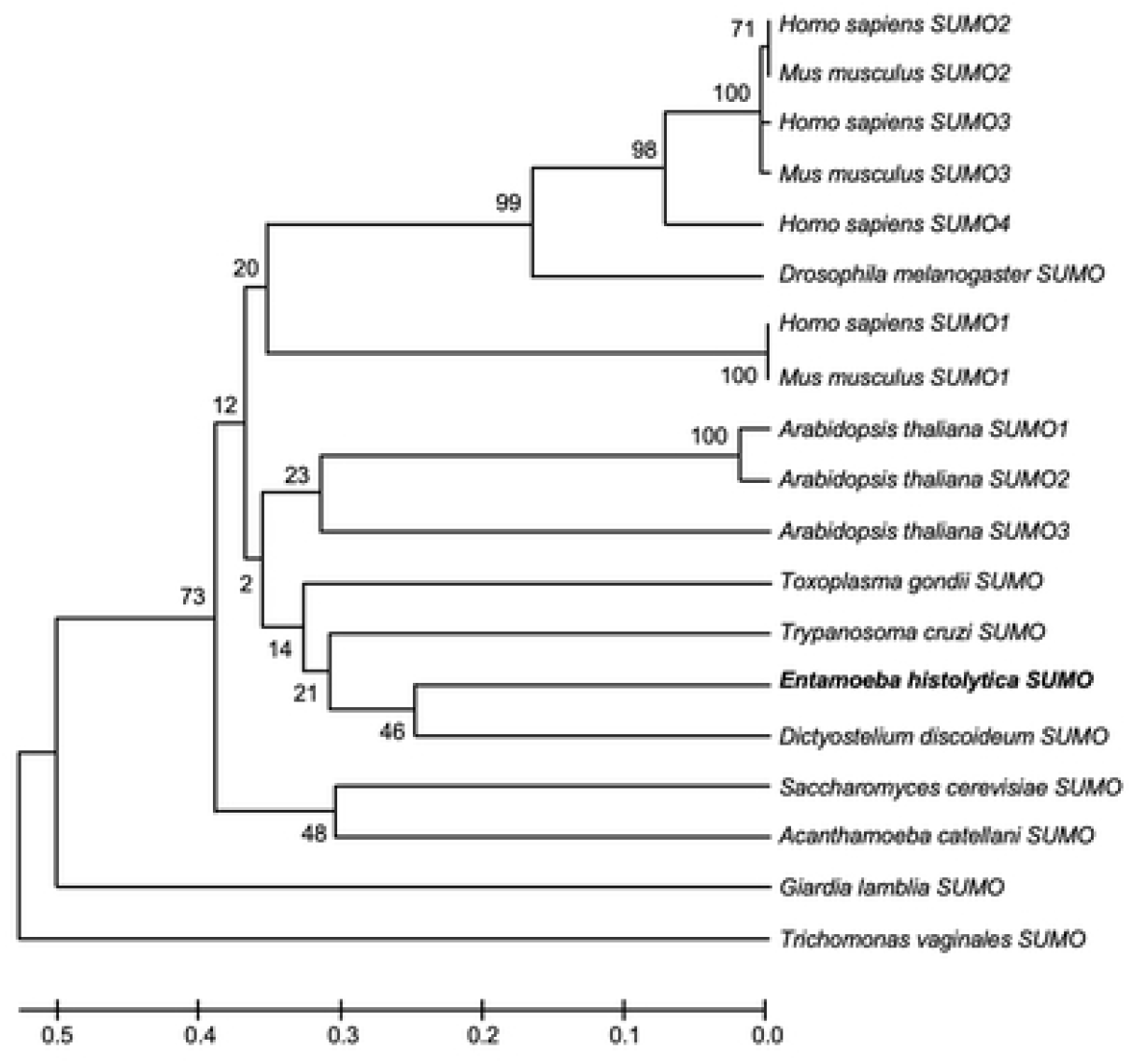
Phylogenetic analysis of EhSUMO. A) Phylogenetic tree performed by UPGMA, using MEGA 5.05 software, shows the position of *E. histolytica* SUMO protein among different species. Numbers in horizontal lines indicate the confidence percentages of the tree topology from bootstrap analysis of 1,000 replicates.

**Table I.**
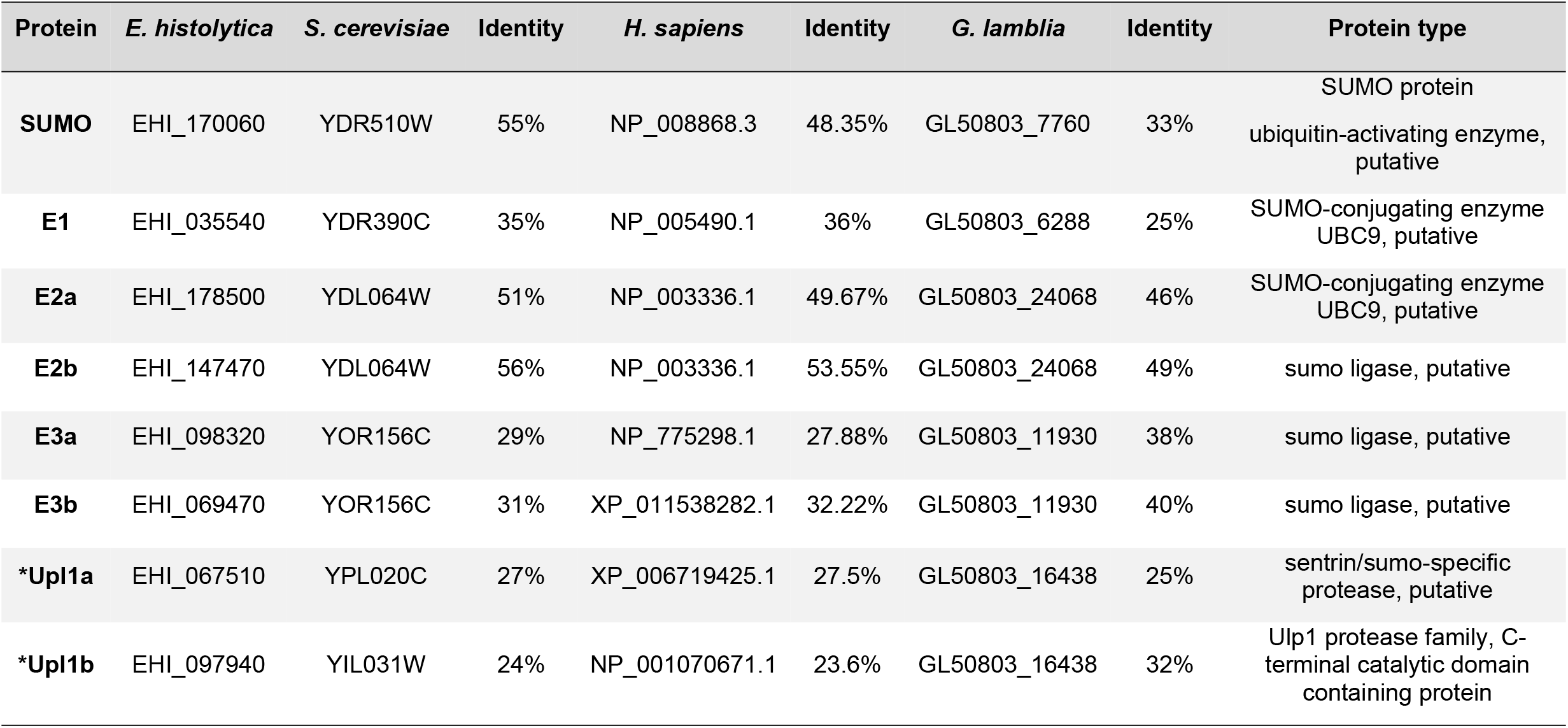
Comparison of putative proteins involved in SUMOylation in *E. histolyitica, S. cerevisiae, H. sapiens* and *G. lamblia*.

**Table II.**
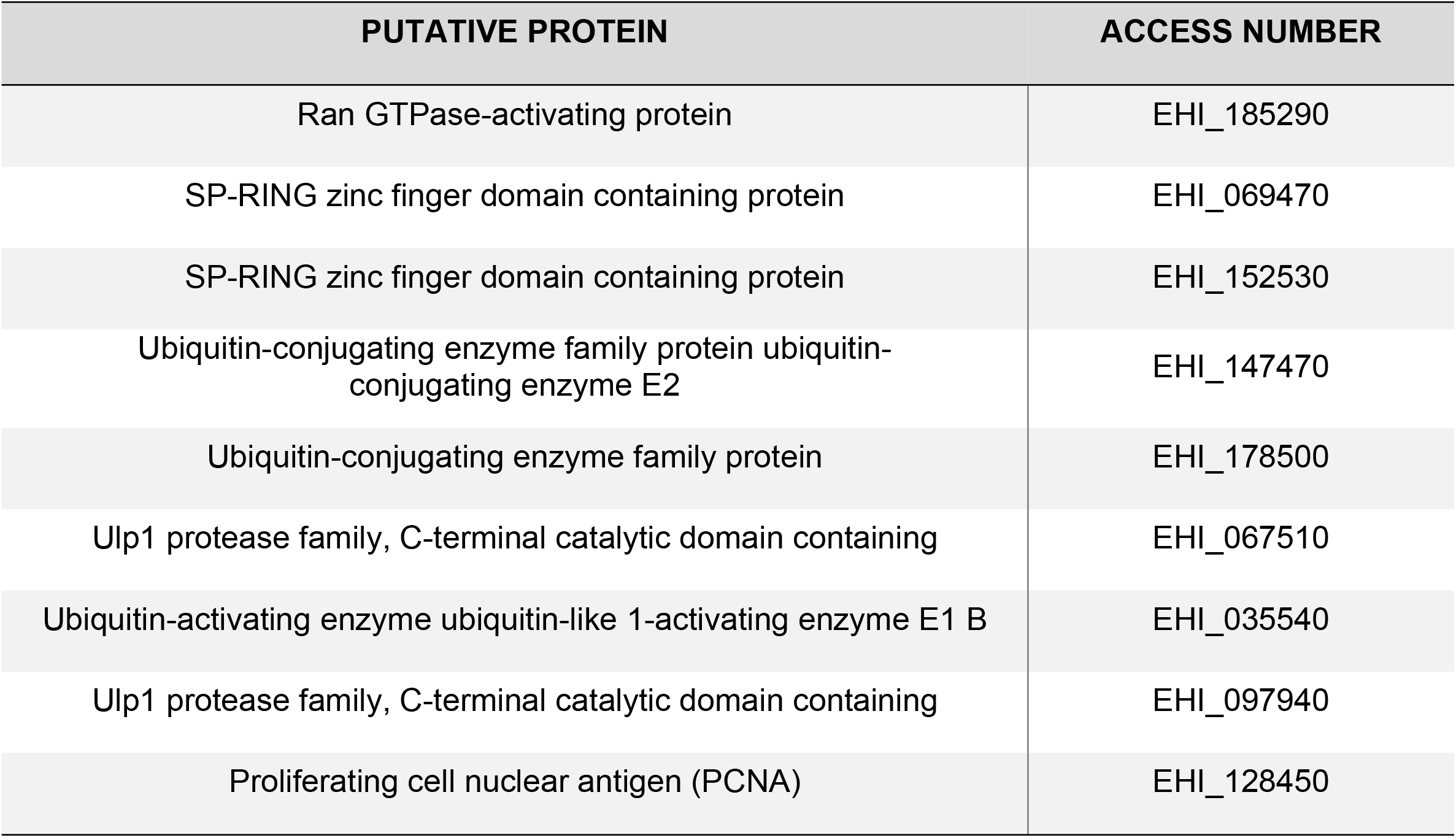
*In silico* interactions of EhSUMO with other *E. histolytica* putative proteins related to the SUMO machinery.

### The 3D-structure of EhSUMO is like other orthologues

To confirm that the sequence that we were working with is a *bona fide* SUMO protein, we obtained the secondary and tertiary structures of the protein. The secondary structure showed the 69 amino acids chain forming the ubiquitin-2Rad60SUMO like-domain (Fig. 3A), with similar size to the one of *G. lamblia* and close to other orthologous (11,19,27–29). The EhSUMO three-dimensional (3D) structure, formed by a single α-helix and four β-strands, overlapped with the ones predicted for G. *lamblia* (RMSD: 0.53), and *S. cerevisiae* (RMSD: 1.26) (PDB:1L2NB), and *H. sapiens* (RMSD:1.20) SUMO-2 (PDB:1A5R) protein crystals (1,30) (Fig. 3B). These findings strengthen the assumption that the EHI_170060 contig corresponds to the phylogenetically conserved SUMO in *E. histolytica*.

**Figure 3.**
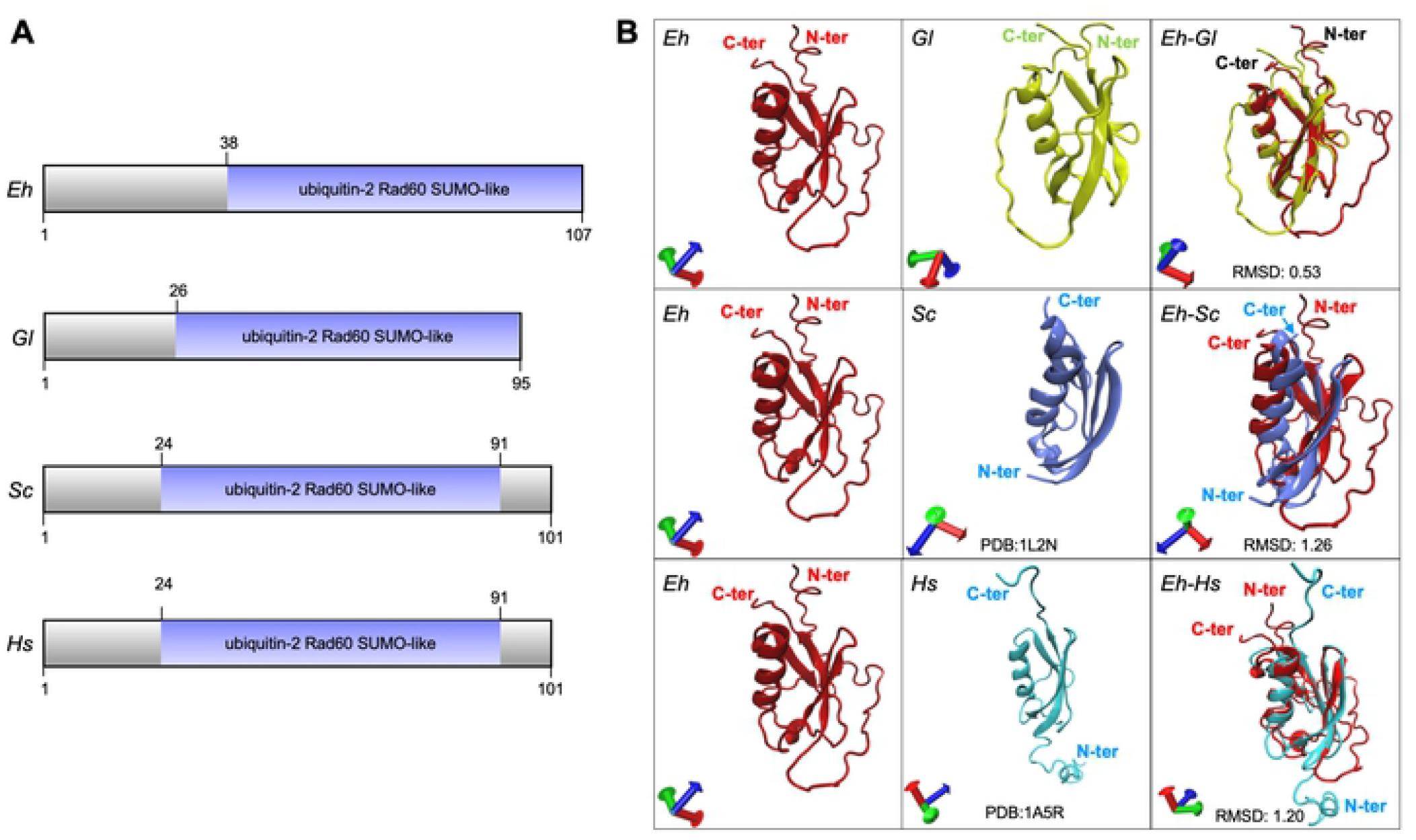
Secondary and tertiary structures of EhSUMO and its orthologous. A) Schematic comparison of functional domains of EhSUMO and other SUMO proteins from different systems. Numbers at the right correspond to the amino acids forming each protein. B) 3D-structures of EhSUMO protein (*Eh*), overlapped with those of *G. lamblia* (*Gl*), *S. cerevisiae* (*Sc*) and *H. sapiens* (*Hs*). The N- and C-terminus regions signaled in red letters. In the bottom of each square are indicated the 3D structures used for *G. lamblia* and *H. sapiens*.

The EhSUMO 3D structure was obtained from the I-TASSER server, after simulations during 200 ns in a soluble environment and then, selected according to its C-score and the best Ramachandran plot values (Fig. 4A). After molecular dynamic simulations (MDS) by the NAMD software, EhSUMO conserved a single α-helix and four β-strands; the rest of the residues appeared lightly twisted in random soft coil and linear structures. The Ramachandran plot showed 98.2% amino acids in the favored regions, 75.7% in the allowed regions, and 1.83% in the outlier ones. Residues distribution indicated that torsion angles of certain amino acids were refined in comparison with those obtained before MDS (Fig. 4A).

RMSD evaluates the system convergence during MDS and indicates whether the values follow a normal distribution (31). EhSUMO reached the equilibrium after 100 ns (Fig. 4B). The Rg values define the protein expansion and compactness. Rg revealed that EhSUMO compacted at the first 100 ns, then, it suffered an expansion from 100 to 135 ns, and in the last 60 ns of the trajectory, EhSUMO again evidenced expansion (Fig. 4C), probably due to the presence of the GG region and its context. Three principal regions appeared as the most flexible areas, detected by RMSF analysis: one at M1 to N35 amino acids composed by coils and turns with non-secondary structure, explaining its higher fluctuation; the second one from Q45 to V50, in a loop, and the last one at the N-terminus from M107 to G114 residues, formed by coils and turns, with a high fluctuation (Fig. 4D). Our findings confirm that EhSUMO has the structure described for other SUMO orthologues.

**Figure 4.**
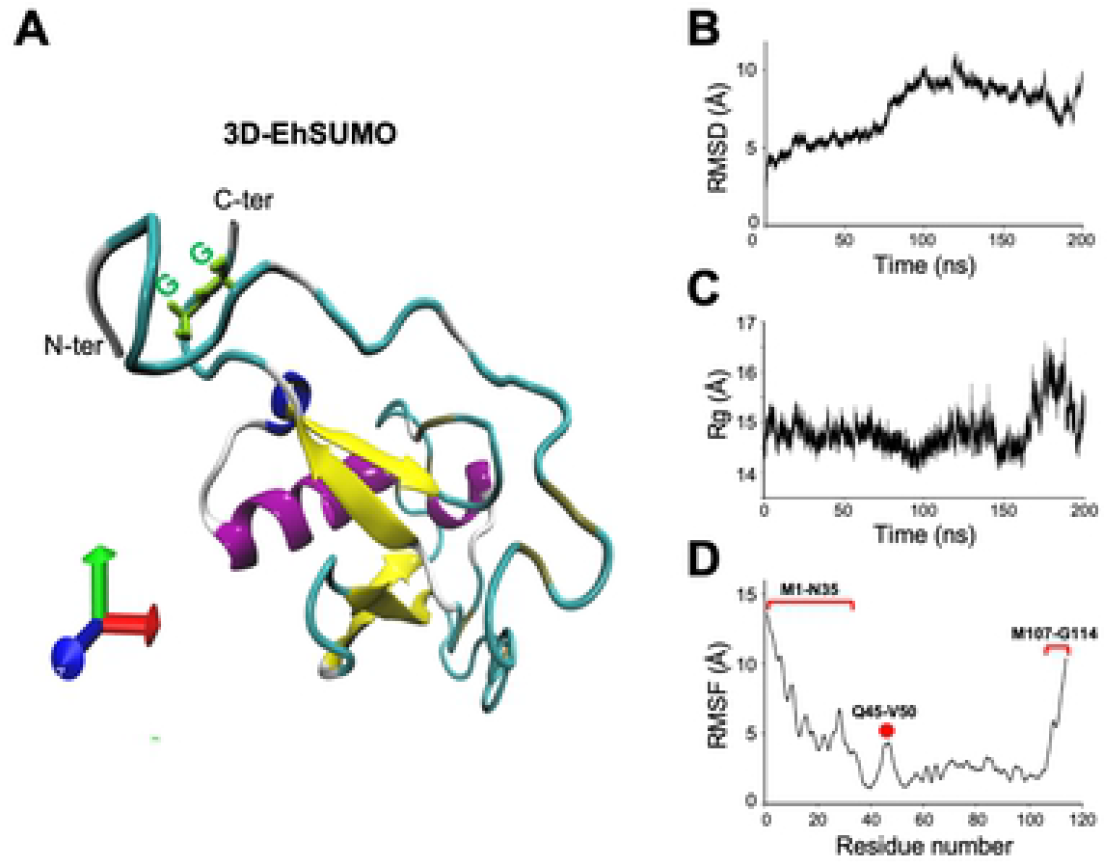
Refined 3D-structure of EhSUMO and MDS. A) Model of EhSUMO presenting the best C-score after 200 ns of MDS. The N- and C-terminus regions, as well as the GG residues are indicated. B-D) The structural analysis of MDS was carried out by RMSD (B), radius of gyration (C) and RMSF (D). The most flexible regions are indicated in red dot and brackets.

### Under the stimulus of erythrocytes, EhSUMO moves from the cytoplasm to the target cell adherence points, phagocytic cups, and phagosomes

According to the *in silico* data, *EhSUMO* could be a *bona fide* orthologous of *SUMO* genes, thus, we proceeded to clone the full gene and express it in *Escherichia coli*. After purification, the recombinant protein (rEhSUMO) was used to obtain rat α-EhSUMO polyclonal antibodies that in western blot assays detected a 17 kDa band in agreement to the 12.6 kDa predicted for EhSUMO plus the histidine label (Fig. 5A).

**Figure 5.**
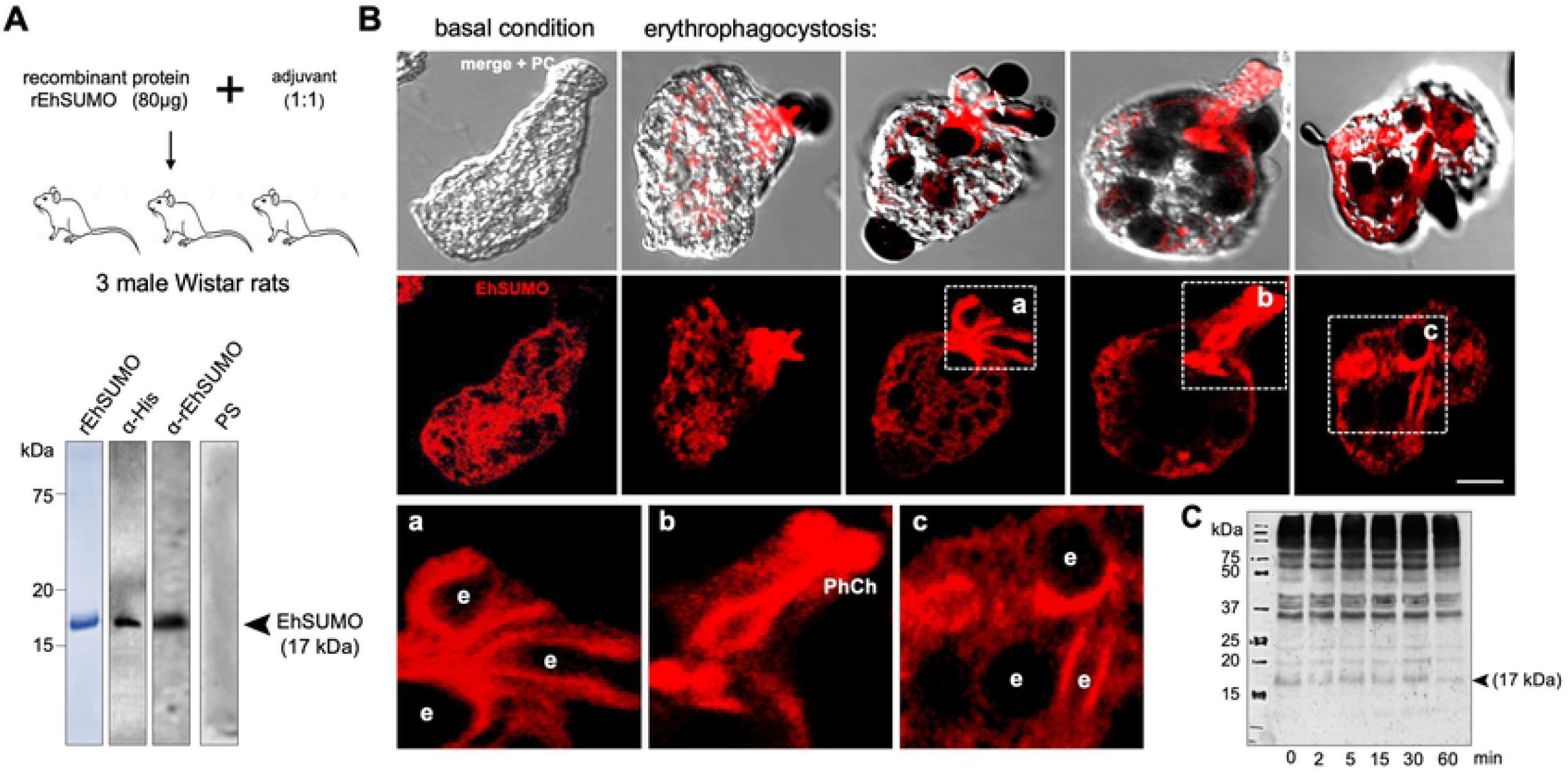
Cellular location of EhSUMO during phagocytosis. A) Immunization scheme to produce α-EhSUMO antibodies, probed in western blot assays, using bacterial lysates. control and pre-immune serum (PS) as Lane 1: rEhSUMO stained with Coomassie blue. Numbers at left: molecular weight standards. B) Confocal microscopy of trophozoites in basal condition and after erythrophagocytosis, using α-EhSUMO antibodies. Squares: different structures formed during phagocytosis magnified in (a-c). a) Structure with form of a conduit or channel. b) Phagocytic channel (PhC). c) Elongated structure surrounding a phagocytosed erythrocyte. e: erythrocytes. Scale bar = 10 μm. C) Western blot analysis of trophozoites lysate after erythrophagocytosis. Number at left: molecular weight standards.

Under confocal microscopy analysis, EhSUMO was located by specific antibodies and fluorescein-labeled rabbit α-rat secondary antibodies. In basal conditions, EhSUMO appeared dispersed in the cytoplasm, close to the internal plasmatic membrane and around vesicles/vacuoles, probably free in the cytoplasm, conjugated or non-conjugated to other molecules, or both (Fig. 5B). After the erythrocytes stimulus was given, EhSUMO moved to the pole where the trophozoites contacted the prey; and fluorescence was more intense in the recently molded phagocytic channels. Fluorescence was also found around the ingested erythrocytes, and in large phagosomes containing three or more erythrocytes (Fig. 5B).

Magnification of these structures revealed the bizarre figures formed in plasma and internal membranes during phagocytosis (Fig. 5B, a-c). However, western blot assays of samples obtained after different times of erythrophagocytosis did not reveal changes in the quality and quantity of proteins. The α-EhSUMO antibodies recognized at least 10 bands from around 17 to more than 240 kDa. Some of these bands might include more than one target protein or contain SUMO polymers (Fig. 5C). The faint band of around 17 kDa could correspond to unconjugated EhSUMO, although Vranych and Merino (19) reported that free SUMO appeared in *G. lamblia* with higher molecular weight than the predicted one, which has been confirmed in several systems (32,33). Intriguingly, the purified recombinant EhSUMO protein migrates at the predicted molecular weight, suggesting that inside the cell, there are other factors that could alter its structure or migration. SUMOylation and deSUMOylation are dynamic events, and the detection of small differences through the phagocytosis kinetics might be hard. Despite this, our results evidence that under the erythrocyte stimulus, EhSUMO moves, together with certain proteins, from the cytoplasm to the phagocytic pole and vesicles, suggesting that SUMOylation could be a switch that prepares proteins to perform their role through phagocytosis.

### *In silico* analysis reveals SUMOylation sites in ESCRT-III and EhADH proteins

One of the long-term goals of our research group has been to discover events governing phagocytosis in *E. histolytica*. Therefore, we investigated if the ESCRT-III proteins and EhADH (17,34), possess sequences that make them susceptible to be SUMOylated. We employed the GPS-SUMO software (35) that predicts attraction sites for SUMO in proteins by an algorithm obtained from 983 SUMOylation sites in 545 proteins and 151 SUMO interaction motifs (SIMs) present in 80 proteins (36). GPS-SUMO software detected putative SUMOylation sites in EhVps2, EhVps24, and EhADH sequences (T**K**LP, V**K**NE, Q**K**AA, respectively) (Table III). Although only EhVps24 conserves the canonical SUMO-binding sequence, the three proteins have the K in the right position. In addition, according to the software, Vps20, EhVps32, and EhADH have SIMs (VTDLDQK IVDLD RQIRQNI, NNEKSHE IGDLL GEDLQDI,EYNSKAQ VILND SKKCES, respectively) (Table III). SIMs facilitate their non-covalent conjugation to SUMO increasing the capacity for SUMOylation, altering the target protein surface and allowing its interaction with distinct molecules (37). These *in silico* results, predict that EhADH and ESCRT-III members are susceptible for being SUMOylated.

**Table III.**
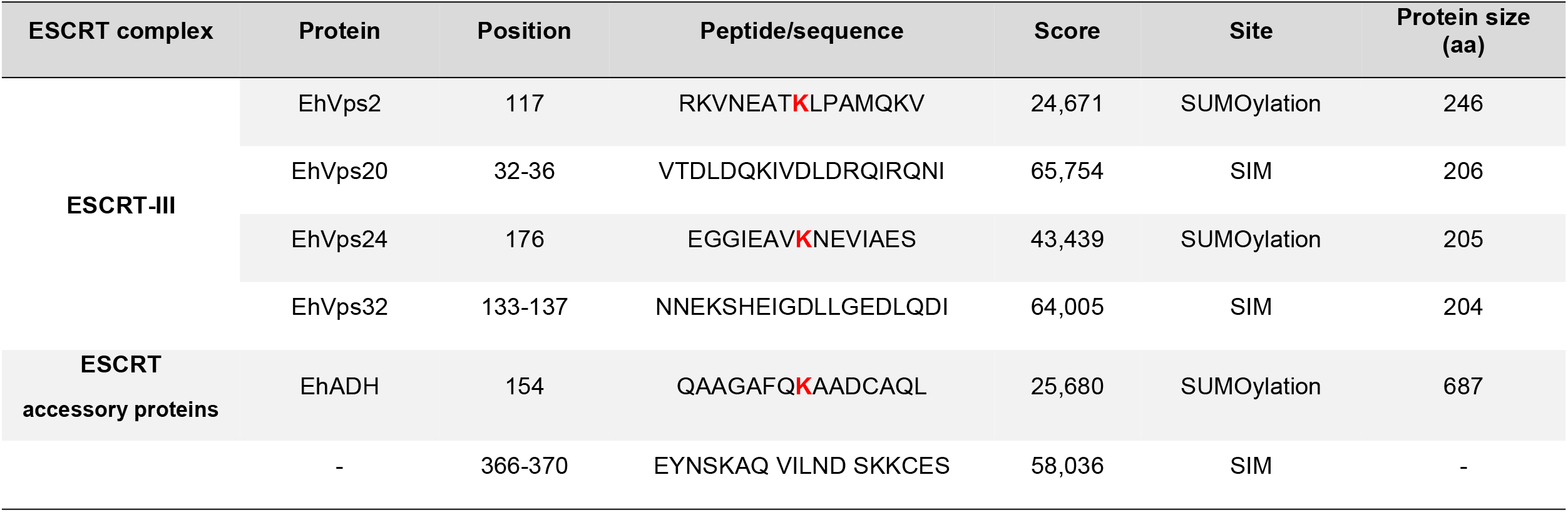
Predicted SUMOylation sites of ESCRT-III and EhADH proteins.

### EhADH protein interacts with EhSUMO

In addition to the SUMOylation sites deciphered by the GPS-SUMO software (35), the secondary structure of EhADH unveiled putative SUMOylation sites at its Bro1 domain (154 amino acid) and at the linker region, from 366 to 370 residues (Fig. 6A). Docking analysis using the 3D model of EhSUMO, obtained here, and the 3D of EhADH previously published (31), suggested that EhADH interacts with EhSUMO by an R rich region, close to the predicted SIM site, making contact also through the S660 to V662 residues, at the C-terminus, whereas EhSUMO interacted with EhADH mainly through the N-terminus with a ΔG = −833.4 (Fig. 6B).

**Figure 6.**
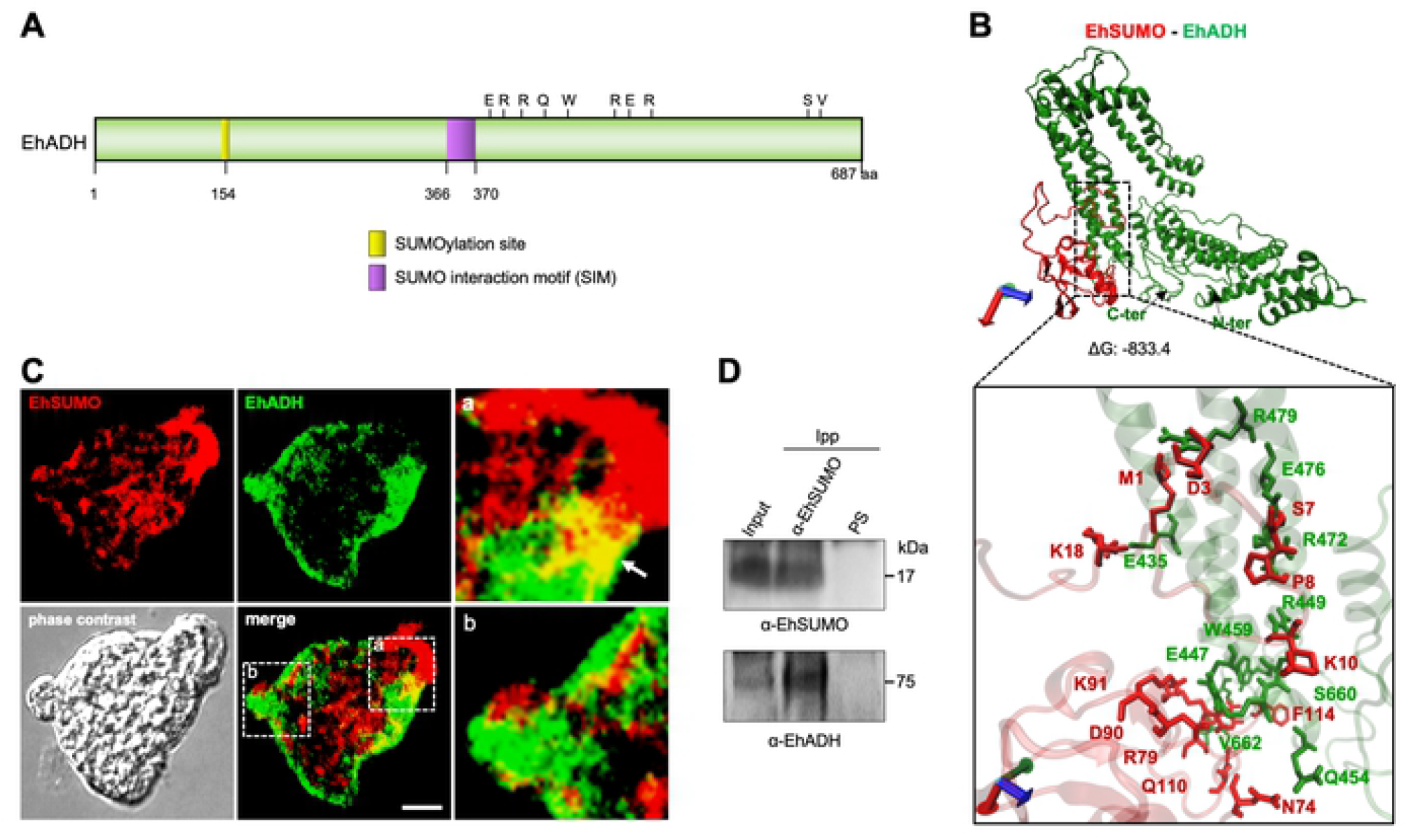
Association of EhSUMO and EhADH. A) Schematic representation of EhADH protein with SUMOylation (yellow) sites and SIM sites (purple). B) Molecular docking between EhSUMO (red) and EhADH (green). Square in the bottom: magnification of the interacting residues. Axes: x, red; y, green; z, blue. C) Immunofluorescence assays of trophozoites under basal conditions using α-EhSUMO (red) and α-EhADH (green) antibodies. Regions in squares are magnified. Arrow: colocalization area. Scale bar = 10 µm. D) Wstern blot of Immunoprecipitates using trophozoites lysates and α-EhSUMO antibody or pre-immune serum (PS) and probed with α-EhSUMO or α-EhADH antibodies. Numbers at right: molecular weight of immunodetected proteins.

Immunofluorescence assays using α-EhSUMO and α-EhADH antibodies, uncovered, in basal conditions, dissimilar fluorescent patterns of EhADH and EhSUMO. However, merging images revealed colocation of both proteins at pseudopodia and in limited regions close to the plasma membrane (Fig. 6C). Immunoprecipitation assays using α-EhSUMO antibodies and trophozoites lysates, confirmed this association. By western blot assays, α-EhSUMO antibodies revealed the EhSUMO protein in the input (total trophozoites proteins), and in immunoprecipitates. Similarly, the α-EhADH antibodies unveiled the EhADH protein in both samples (Fig. 6D). These results strongly suggest that both proteins associate each other, interacting directly or indirectly.

### Colocation of EhSUMO and EhADH increases during phagocytosis

To investigate the cellular fate of EhSUMO and EhADH during phagocytosis, trophozoites were prepared for immunofluorescence assays and examined through the confocal microscopy after different times of phagocytosis. At 5 min, EhSUMO was detected in the place of contact where erythrocytes were being ingested (Fig. 7A, B), forming the peculiar membranous structures in the phagocytic channel (see Fig. 5). It also appeared around ingested erythrocytes and surrounding big pockets that could be the pre-phagosomes formed to receive the red blood cells from other endosomes (Fig. 7A). In some images, EhADH did not show up in the same region of the phagocytic channel that EhSUMO did, although laser sections revealed its presence in other places of this structure, as published before (38). Large bags in the cytoplasm, that could correspond to a putative pre-phagosome structures, were also enlightened by the α-EhADH antibodies, inside and surrounding them, and the label was also detected around the ingested erythrocytes (Fig. 7A). At 15 and 30 min of phagocytosis, the EhADH and EhSUMO colocation increased (Fig. 7), and at 60 min, both proteins moved to the internal plasma membrane and remained around the ingested erythrocytes. The erythrocytes-containing structures exhibited different size and number of erythrocytes inside. Some of them, emerged stained only by α-EhSUMO or by α-EhADH antibodies, suggesting a differential participation for SUMOylated proteins during the maturation of endosomes/phagosomes. The three conditions were observed: EhSUMO and EhADH separated and both proteins associated. Quantification of fluorescence colocation confirmed that it increased through the phagocytosis kinetics (Fig. 7C), suggesting that the EhSUMO-EhADH association enhance during this virulence event.

**Figure 7.**
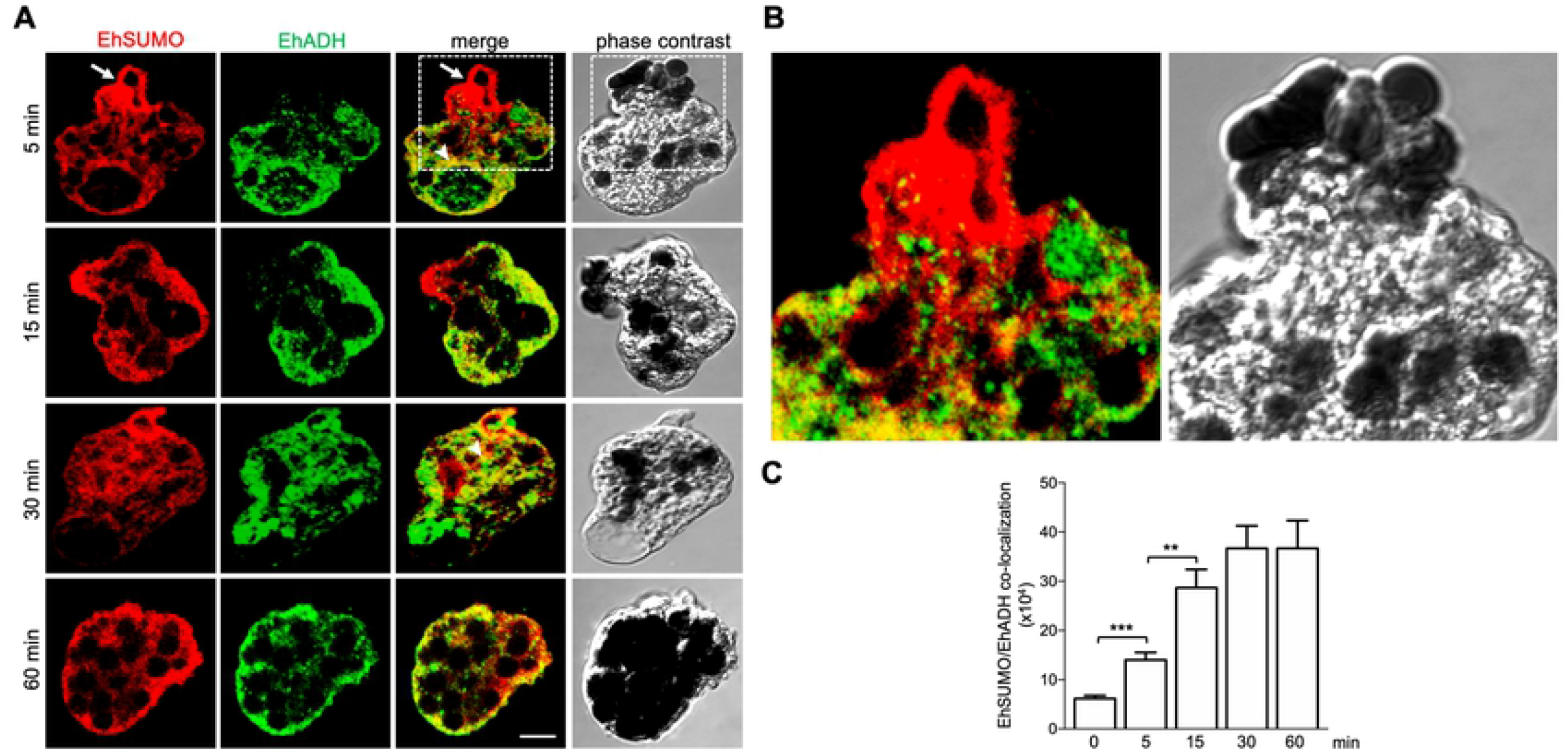
Location of EhSUMO and EhADH during phagocytosis. Trophozoites were incubated for 5, 15, 30 and 60 min with erythrocytes and processed for confocal microscopy. A) Immunofluorescence assays using α-EhSUMO (red) and α-EhADH (green) antibodies. Arrow: EhSUMO in erythrocytes being phagocytosed. Scale bar = 10 µm. B) Magnification of squares in (A). C) Quantification of EhSUMO and EhADH colocalization in the whole cells. (**) *p* < 0.01 (***) *p* < 0.001.

### *In silico* analysis discloses interaction between EhVps32 and EhSUMO

EhVps32 has a predicted molecular weight of 22.4 kDa, although due to its structure and charge of amino acids, it migrates at 32 kDa in SDS-PAGE (14). Its secondary structure exhibits a SIM (NNEKSHE IGDLL GEDLQDI) from the 134 to 137 amino acids (Fig. 8A), whereas its 3D-structure, obtained from the I-TASSER server, presents three α-helices, after 200 ns MDS by NAMD software in a soluble environment (Fig. 8B). The model selected according to the C-score and the best Ramachandran plot values was employed for these analyses.

**Figure 8.**
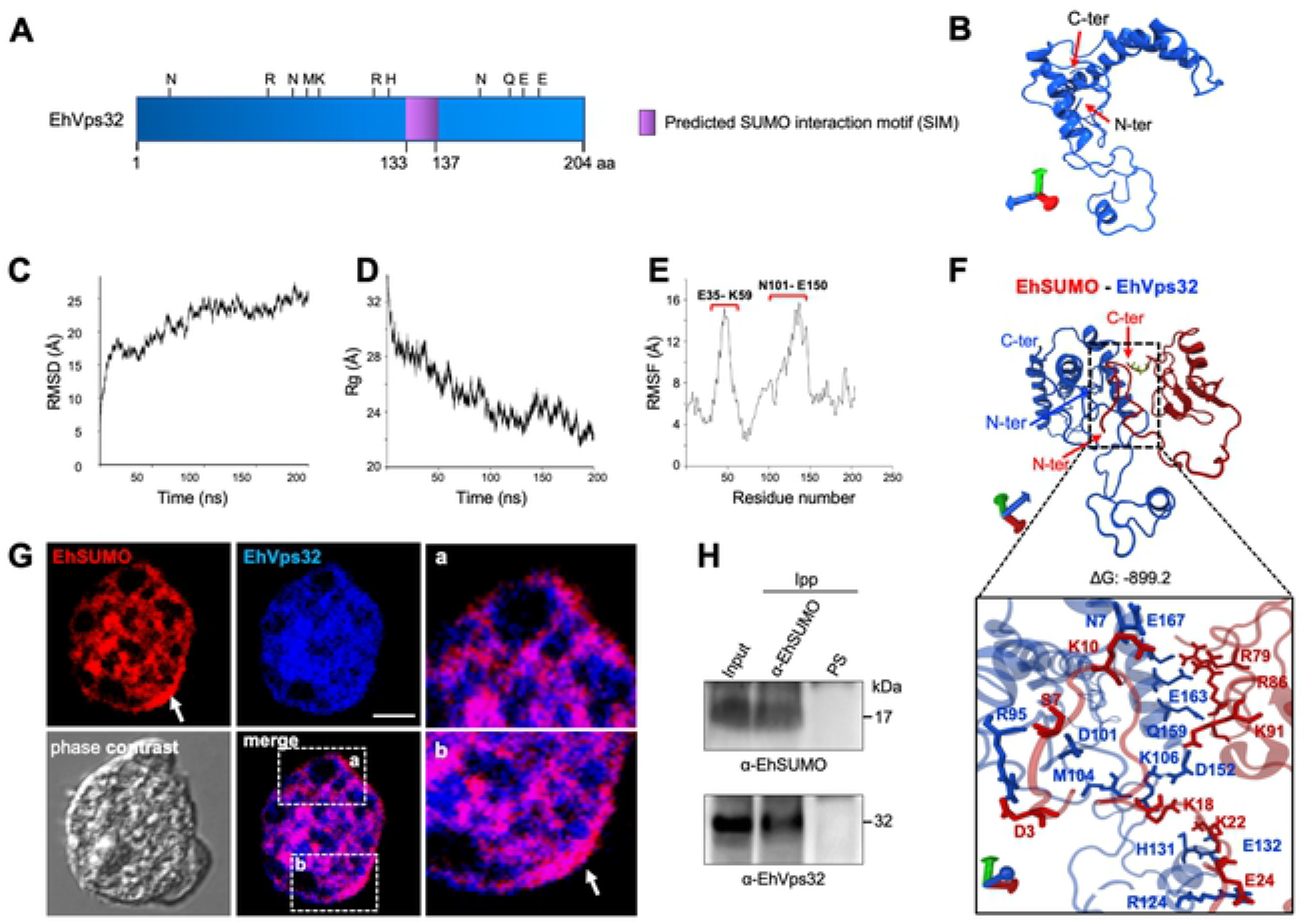
Association of EhSUMO and EhVps32. A) Schematic representation of SIM (purple) site in EhVps32 protein. B) Model of EhVps32 presenting the best C-score after 200 ns of MDS. The N- and C-terminus regions are indicated. C): Structural analysis of MDS carried out by RMSD, D): radius of gyration, E): RMSF Red brackets indicate the most flexible regions. F) Molecular docking of EhSUMO (red) and EhVps32 (blue). Square: magnification of the interacting residues. Sticks in green indicates the glycines in EhSUMO. Axes: x, red; y, green; z, blue. G) Immunofluorescence assays of trophozoites under basal conditions using α-EhSUMO (red) and α-EhVps32 (blue) antibodies. Squares: magnification of proteins colocalization (a, b and arrow). Arrow: area of colocalization in the internal plasma membrane. Scale bar = 10 µm. H) Trophozoites in basal conditions were lysed and immunoprecipitated using α-EhSUMO antibody or pre-immune serum (PS) and immunoprecipitated proteins were analyzed by western blot using α-EhSUMO or α-EhVps32 antibodies. Numbers at right: molecular weight of immunodetected proteins in western blot assays.

The Ramachandran plot showed 92.98% of the amino acids in the favored regions, 69.47% in the allowed regions, and 3.16% in the outlier ones. Besides, the RMSD analysis showed that EhVps32 reached the equilibrium at 100 ns (Fig. 8C), whereas the Rg values, indicated that it compacted through the trajectory (Fig. 8D) exhibiting two regions as the most flexible areas, located at E35 to K59 amino acids and the between N101-E150 residues, respectively. Docking analysis predicted that EhVps32 and EhSUMO contact each other with a ΔG= −899.24 (Fig. 8F). As in the case of EhADH, the interaction was performed in a wider region than the predicted one, distributed along the whole protein. The site was composed of three N, one close to the amino terminus, other to the C-terminus and the third one at RNMK motif (Fig. 8F), while EhSUMO contacted EhVps32 through the N-terminus (Fig. 8F). These bioinformatics data predict the association between EhSUMO and EhVps32 proteins.

### Immunofluorescence and immunoprecipitation experiments confirm the EhSUMO and EhVps32 interaction

Confocal images of trophozoites in basal conditions, evidenced colocation of EhSUMO and EhVps32 around vesicles/vacuoles, in the inner plasma membrane, in the origin of the pseudopodia and extensively in clumps in the cytoplasm (Fig. 8G). Immunoprecipitation assays, using α-EhSUMO antibodies revealed EhSUMO and EhVps32 in the immunoprecipitates (Fig. 8H). All these experiments showed a strong association between EhSUMO and EhVps32, although it could be direct or indirect.

### Interaction between EhVps32 and EhSUMO continues through phagocytosis

Through phagocytosis, confocal images using the α-EhSUMO antibodies, disclosed the same membranous structures described above, and the small vesicles surrounding larger endosomes/phagosomes (Fig. 9A, B). Whereas the α-EhVps32 antibodies appeared around the erythrocytes, in the phagocytic bags, and close to the internal plasma membrane (Fig. 9A), as described (17). Both proteins exhibited an extensive colocation, suggesting that EhSUMO is associated to EVps32 or to other proteins interacting with EhVps32, which could include the ESCRT-III members and EhADH (Fig. 7). However, red spots appeared around to and near of the red blood cells (Fig. 9), showing that, EhSUMO also conjugates to other proteins that are not in contact with EhVps32. In contrast to the experiments performed with α-EhADH and α-EhSUMO antibodies, colocation of EhVps32 and EhSUMO was stronger, and it was maintained and even slightly increased during phagocytosis (Fig. 9 A, B). Quantification of the fluorescent label in colocation confirmed this (Fig. 9C).

**Figure 9.**
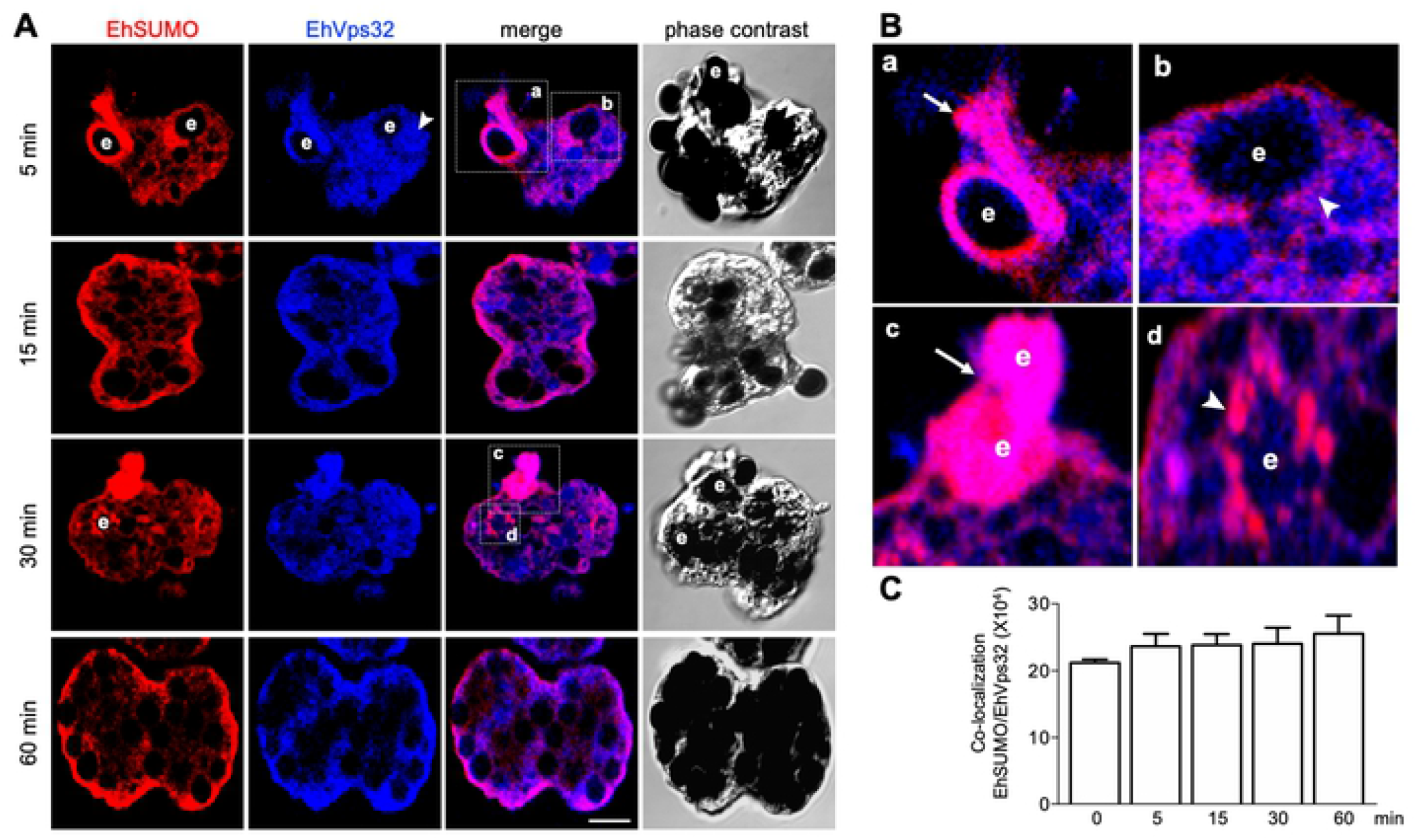
Location of EhSUMO and EhVps32 during phagocytosis. Trophozoites were incubated for 5, 15, 30 and 60 min with erythrocytes (e) and processed for confocal microscopy. A) Immunofluorescence assays, using α-EhSUMO (red) and α-EhVps32 (blue) antibodies. Scale bar = 10 µm. B) Magnification of squares presented in (A). Arrow: colocalization of proteins in a duct-like structure (a) and in erythrocytes (c). Arrowhead: colocalization of both proteins surrounding a vesicle with erythrocytes (b, d). C) Quantification of EhSUMO and EhADH colocalization in the whole cells.

### *EhSUMO* knocked-down trophozoites have a lower rate of erythrophagocytosis than the wild type strain

To deepen the importance of SUMOylation in phagocytosis, we silenced the *EhSUMO* gene, using double-stranded RNA (dsRNA), expressed in bacteria, (39). After incubation of trophozoites for 24 h with the dsRNA, lysates from silenced (KD) and control trophozoites were submitted to western blot assays. Protein patterns of control in KD trophozoites notably differed (Fig. 10A). In the control, bands from about 17 kDa to more than 240 kDa reproducibly appeared in the nitrocellulose membranes (like it is shown in figure 5). However, in the KD trophozoites, bands were fainter, and some proteins of high molecular weight did not appear (Fig. 10 A), suggesting a lower SUMOylation of certain proteins. The antibodies against the nuclear protein EhPCNA, evidenced that the amount of protein used in both lanes of the gel was similar, and degradation was not detected, at least for this protein (Fig. 10 A).

**Figure 10.**
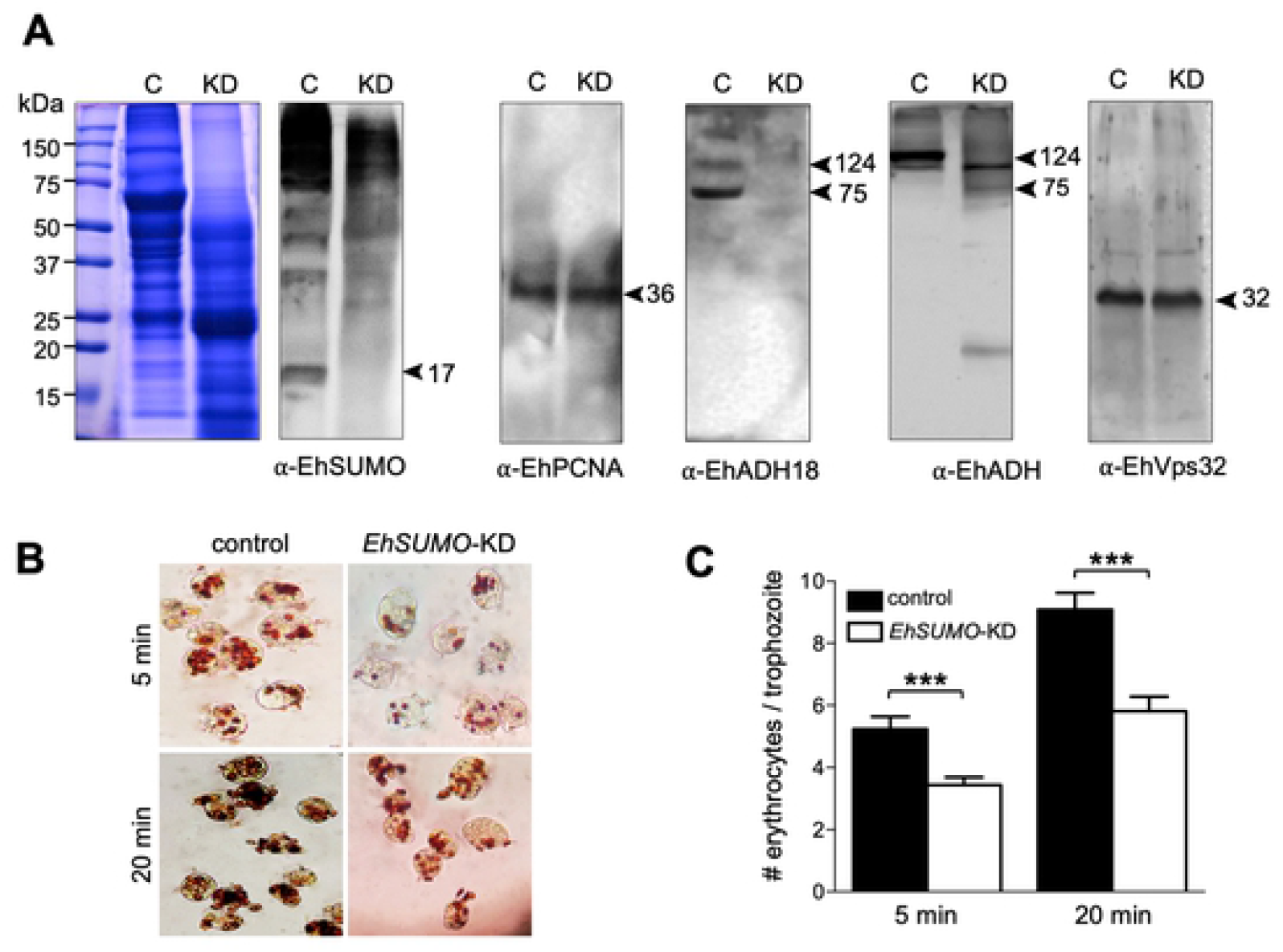
Effect of EhSUMO knock-down in the expression of EhADH and EhVps32 proteins and in phagocytosis. Trophozoites were silenced using the pL4440/*EhSUMO* plasmid as described in materials and methods. A) Western blot of lysates from control (C) or EhSUMO-KD trophozoites (KD) in basal condition, using different antibodies: α-EhSUMO, α-EhADH, α-EhADH-18 or α-EhVps32. α-EhPCNA: loading control. Numbers at left: molecular weight standards. Number at right: molecular weight in kDa of immunodetected proteins. B) Novikoff staining of control and EhSUMO-KD trophozoites at 5 and 20 min erythrophagocytosis. C) Rate of erythrophagocytosis was evaluated by counting the number of erythrocytes per trophozoite. Data represent the media and standard error of 100 trophozoites. (***) *p* < 0.001.

Polyclonal antibodies against an exposed peptide in the EhADH structure (αEhADH18), (494-KFRQFENDIKLLCEGNIQ-513 residues) recognized in the control trophozoites, the 75 kDa band of EhADH and the 112-124 kDa bands representing the distinct conformations of EhCPADH complex. However, only faint bands were detected in KD cells (Fig. 10A). This complex is formed by association of EhADH with the EhCP112 cysteine protease (31,34) and it acquires distinct conformations, presenting slight differences in migration in SDS-PAGE. SUMOylation-deSUMOylation alters protein conformation, hiding relevant epitopes for protein binding and antibody recognition (40). To explore this idea, we used a polyclonal antibody, directed against the full length EhADH protein (α−EhADH). In agreement with earlier results (34,41), in the control trophozoites, α−EhADH antibodies recognized the 112-124 kDa bands corresponding to the EhCPADH complex (41), but free EhADH was not visible. Surprisingly, in KD trophozoites, the α−EhADH antibodies detected the 75 kDa band corresponding to free EhADH and the EhCPADH complex (Fig. 10 A). These results support the hypothesis of alteration of EhADH conformation, probably due to the lack of efficient SUMOylation. However, this assumption needs more experiments to be precisely confirm it. Interestingly, EhVps32 appeared likewise in KD and control trophozoites, also serving as an internal control of the amount and integrity of proteins loaded in the gel. Although we would expect distinct molecular weights between the SUMOylated and non-SUMOylated EhVps32, it is possible that during the sample preparation process, the non-covalent binding of SUMO to its target could be destroyed, or that EhSUMO could be indirectly bound to EhVps32. Erythrophagocytosis experiments evidenced that in all cases, KD trophozoites phagocytosed between 25 and 30% fewer erythrocytes than the wild type (Fig. 10 B, C).

Laser confocal immunofluorescence experiments verified the effect of *EhSUMO* silencing in trophozoites. The α-EhSUMO antibodies recognized stronger the control trophozoites than the KD ones. Additionally, the bizarre membranous structures formed by α-EhSUMO antibodies were not visible in KD trophozoites during phagocytosis. Besides, in contrast to the western blot assays, α-EhADH and α-EhVps32 antibodies were less reactive with KD trophozoites in immunofluorescence assays (Fig. 11A). The three proteins appeared associated in different areas in the control, even in basal conditions, but in KD trophozoites, colocation was much lower (Fig. 11B, C). During erythrophagocytosis, EhSUMO-deficient trophozoites displayed a poor colocalization and a significant reduction of the recognition of EhADH and EhVps32 by the antibodies (Fig. 12). These findings demonstrate the relevance of SUMOylation in phagocytosis and evidence that proteins are affected in distinct ways by the SUMOylation-deSUMOylation processes.

**Figure 11.**
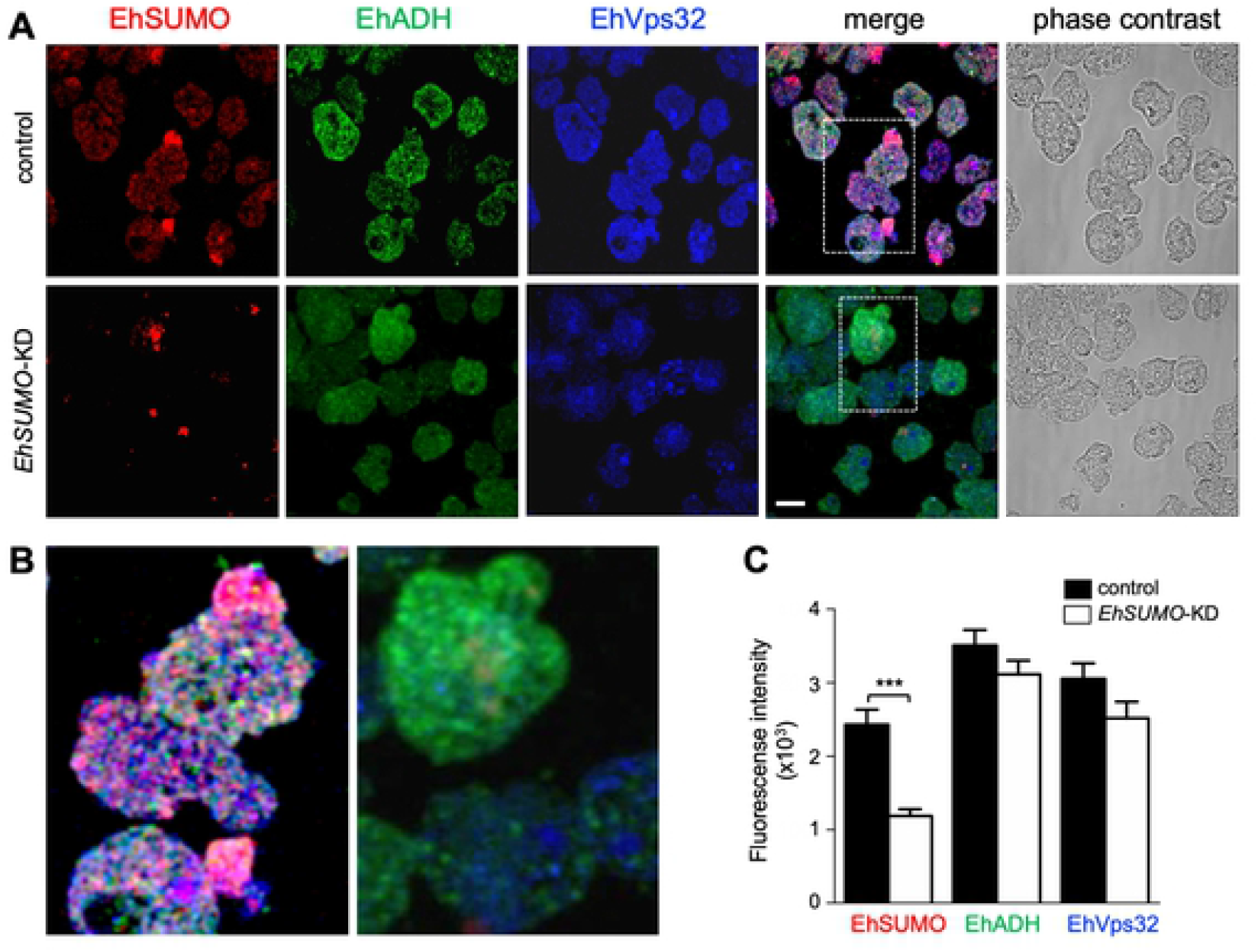
Localization of EhSUMO, EhADH and EhVps32 in EhSUMO-KD trophozoites in basal conditions. Confocal microscopy of control and KD trophozoites in basal conditions A) Representative image of control and EhSUMO-KD trophozoites using α-EhSUMO (red), α-EhADH (green) and α-EhVps32 (blue) antibodies. Scale bar = 10 µm. B) Magnification of squares in (A). C) Fluorescence intensity measured by pixels and corresponding to EhSUMO, EhADH and EhVps32 in both types of trophozoites. (***) *p* < 0.001.

**Figure 12.**
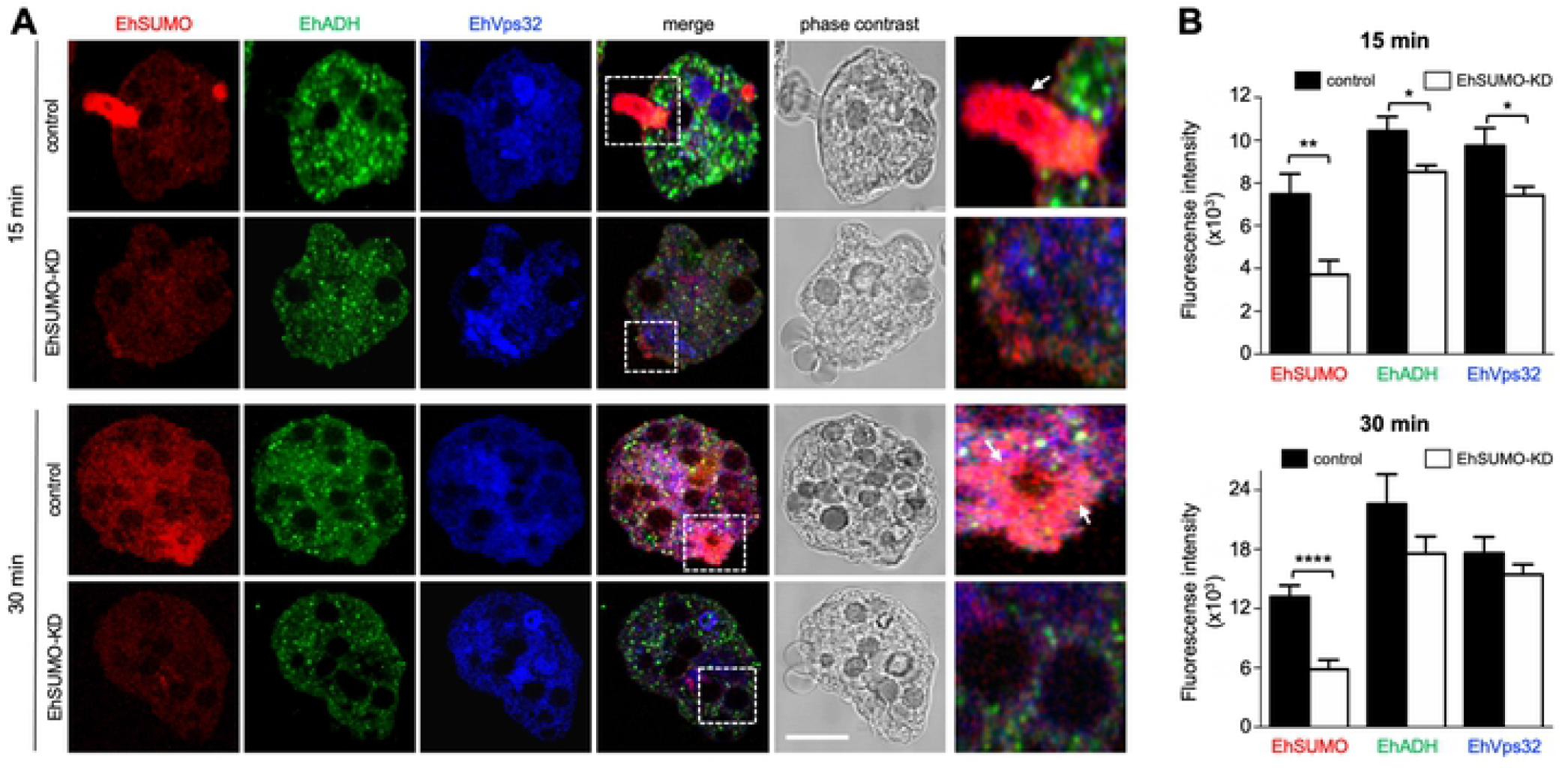
Localization of EhSUMO, EhADH and EhVps32 in EhSUMO-KD trophozoites during phagocytosis. Erythrophagocytosis kinetics of trophozoites silenced using the PL4440/*Ehsumo* plasmid. A) Representative image of control and EhSUMO-KD trophozoites at 15 and 30 min of phagocytosis using α-EhSUMO (red), α-EhADH (green) and α-EhVps32 (blue) antibodies and analyzed by confocal microscopy. Scale bar = 10 µm. Right panels: Magnification of squares in merging images. Arrows: phagocytic channel and areas around erythrocytes B) Fluorescence intensity measured by pixels and corresponding to EhSUMO, EhADH and EhVps32 proteins in both types of trophozoites at both time of phagocytosis. (*) *p* < 0.05, (**) *p* < 0.01 and (****) *p* < 0.0001.

In conclusion, altogether, the results presented here prove the presence of an intronless *bona fide EhSUMO* gene encoding for a predicted 12.6 kDa protein that actively participates in phagocytosis. Silencing of the *EhSUMO* gene affected the rate of phagocytosis and possibly the EhADH and EhVps32 function, supporting the importance of SUMOylation in phagocytosis, a landmark for the parasite virulence.

## Discussion

In this paper, we disclosed and characterized the presence of a *SUMO* gene and its product in *E. histolytica*. Then, by bioinformatic analysis, we also found in the AmoebaDB the enzymes required for SUMOylation of target proteins: E1, the activating enzyme, E2, the conjugated one, and E3 the ligase, as well as those participating in deSUMOylation: UIpb1a and UIpb1b. Our data strongly suggest that SUMOylation is a modifier of the parasite proteins, stimulating some of them for phagocytosis. Furthermore, we unveiled the association of EhSUMO, a ubiquitin-like modifier (UbI) protein, with EhADH and EhVps32 proteins, both involved in phagocytosis. Beyond the detection and characterization of EhSUMO in this parasite, the relevance of this work relies on two main aspects: i) this is the first report on the role of SUMOylation in phagocytosis and in the modification of specific proteins participating in this phenomenon, and ii) the involvement of SUMOylation during phagocytosis highlights the potential use of this knowledge for the development of therapeutics and diagnosis methods to defeat amoebiasis.

Lack of vaccines, reduced chemotherapy options, and the emergence of drug-resistant parasites (42) are challenges presented by diseases caused by protozoa, among these, amoebiasis. PTMs, including UbIs, modify proteins to facilitate their functions, including virulence-related functions (43). UbIs have been widely studied in yeast, plants, and Mammals (43,44), but little is known on their role in protozoan parasites, highly divergent organisms with characteristics that might be investigated to understand their evolutionary process and their virulence mechanisms (45,46). Ubls impact the regulation of cellular functions such as cell cycle progression (11), transcription (47,48), stress responses (49), DNA damage repair (50), cell signaling (22), nuclear transport (51), and autophagy (43). By findings reported here, we add to this list, an old event, found now to be regulated by SUMO: the phagocytosis, involved in damage produced by *E. histolytica* trophozoites.

In *E. histolytica*, Arya et al. in 2012 (52) reported a bioinformatic study on UbIs modifiers and their conjugated enzymes, and recently, Kumari et al. In 2018 (53) found a UBc7/Ube2g2 protein connected with the plasma membrane and phagocytosis in trophozoites. Beside this, we have not found reports on SUMOylation in this parasite. By *in silico* analysis, we detected EHI_170060 contig that presents the characteristics described so far for SUMO. In the phylogenetic tree, EHI_1700060 product (*EhSUMO*) appeared close to *T. cruzi* and *D. discoideum* orthologues. However, we used here as a template the yeast and human SUMO-2 proteins, because they have been extensively studied, their 3D structures are well known, and crystal are available. HsSUMO-2 forms stable polymeric chains that also are susceptible to poly-ubiquitination, a signal for proteasome degradation (54). This effect is given by the K11 of the ΨKxE/D consensus motif that allows the formation of poly-SUMO chains, absent in HsSUMO-1. Although we have not studied its relevance of this K in trophozoites, EhSUMO possesses the K amino acid at the motif ΨKxE/D, but at K22. These facts and others, including the MDS analysis (1,55), support the identity and nature EhSUMO

Docking of EhADH and EhSUMO, and EhVps32 and EhSUMO, predicted that EhSUMO uses different motifs in the protein target to join to other proteins, but targets suggested by our results could need fine tuning of punctual mutations to precisely determine the binding sites. However, the confocal images evidenced that EhADH and EhVps32 efficiently bind, directly or indirectly, to EhSUMO, and that SUMOylation influences their cellular location, although we cannot discern how much of the changes were due to the phagocytosis event and how much to the SUMOylation process. Nevertheless, the fact that KD trophozoites in basal conditions, did not display EhADH close to the plasma membrane or in pseudopodia, suggests that EhADH need to be SUMOylated to find its cellular position. Confocal assays using control and KD trophozoites, support this assumption. Altogether, these results point out to the hypothesis that the equilibrium between PTM proteins and their unmodified state modulates the fate and function of the target substrates during phagocytosis, granting the fine-tuning of the cellular mechanisms needed for life.

Except for EhADH and EhVps32, we could not disclose changes in the amount and nature of the proteins that are SUMOylated during phagocytosis, but SUMOylation-deSUMOylation is a highly dynamic process and undetectable changes could be occurring. The multiple and speedy changes produced during this phenomenon were evidenced by the images obtained using α-EhSUMO antibodies that uncovered the active participation of EhSUMO. The images revealed the extensive membrane changes accordingly to the moment of erythrocyte’s contact and ingestion during the intense vesicular trafficking accelerated by phagocytosis. In fact, we did not find a single repetitive bizarre figure in a trophozoite or in the total population, corroborating the dynamics of the event.

SUMOylation alters not only the cellular location of the target protein, but changes the protein conformation, and, consequently, their affinity to other proteins (56). This has been studied mainly in mammalian cells and yeast, but little in parasites, even when these events might be part of their virulence mechanisms. The change of EhADH conformation was suggested by the lack of recognition by the α-EhADH18 antibodies in KD trophozoites. To discard protein degradation, we used a polyclonal antibody directed to the full-length EhADH protein that detected the complex in both types of trophozoites and reacted with the 75 kDa band, evidencing the integrity of the protein. We suspect that the SUMOylation could produce stability on EhADH, but when EhSUMO is diminished, the non-SUMOylated protein could change its structure. In fact, MDS experiments characterizing the different protein conformations, showed that EhADH assumes distinct structures (31).

The position of EhADH and EhVps32 inside the cell, after they were hypothetically SUMOylated (deduced by the colocation images), constantly change during phagocytosis. Each protein behaved in a different way: colocalization of EhADH and EhSUMO significantly increased through phagocytosis, whereas EhVps32 and EhSUMO maintains a high level of colocalization since basal state. These differences could be interpreted as necessary events to carry out the distinct functions of the proteins. Our work with EhADH since many years ago (41), has evidenced its interaction with many other molecules, such as EhVps32 (14) and EhNPC1 and EhNPC2 (57) as it has been described for other ALIX family proteins (58), thus, it is possible that the protein conformation could be altered according to the target protein. Meanwhile, the EhVps32 movement is probably limited to the ESCRT-III complex participation. Hence, SUMOylation could enable them to move in the cell and perform their functions.

Karpiyevich and Artavanis-Tsakonas in 2020 (44) postulated that in addition to fulfilling the conserved functions described for these modifiers, Ub1s participate in novel parasite-specific roles. The data presented here, support the Karpiyevich and Artavanis-Tsakonas hypothesis (44), suggesting that proteins involved in phagocytosis of *E. histolytica* trophozoites suffer SUMOylation, as a requisite to carry out their tasks. In conclusion, our findings point out the importance of SUMOylation in phagocytosis. Based on these data, it is possible that in the future, inhibition of SUMOylation in *E. histolytica* and in other parasites could help to find novel therapeutic methods to defeat amoebiasis and other parasitic diseases.

### Materials and methods Culture of trophozoites

Trophozoites of *E. histolytica*, clone A (strain HM1: IMSS) (Orozco et al., 1983), were axenically cultured in TYI-S-33 medium at 37*°*C and harvested in logarithmic growth phase for all experiments (59).

### SUMO searching and phylogenetic analysis

Full-length sequence of *SUMO* gene was identified in the *E. histolytica* genome, using the BLASTP algorithm (http://blast.ncbi.nlm.nih.gov//Blast.cgi) and the AmoebaDB database (www.amoebadb.org). Additionally, SUMO proteins of *G*.*lamblia* (accession number GL50803_7760), *H. sapiens* (accession number NP_001005781) and *S. cerevisiae* (accession number NP_010798.1), were also explored. The *EhSUMO* sequence was analyzed and 5’ and 3’ en primers were designed, to amplify the full-length gene. The putative amino acid sequence of EhSUMO (EHI_170060) was aligned with its orthologous by ClustalW and data were submitted to phylogenetic analysis by UPGMA, using MEGA 5.05 software. The bootstrapping was performed in 1000 replicates.

### Secondary and tertiary structure of EhSUMO

By *in silico* analysis, the ubiquitin-2 Rad60 domain, characteristic of SUMO proteins, was located and aligned in the putative EhSUMO amino acid sequence and compared with SUMO orthologous, to design the secondary structure of the protein. The 3D structure of EhSUMO was obtained using the crystal of *H. sapiens* (PDB:1A5R) and *S. cerevisiae* (PDB:1L2N). For *G. lamblia* SUMO 3D-structure, we submitted the protein sequence V6TGL retrieved from UniProtKB, to I-TASSER server and the higher C-score was selected. The alignment of proteins was visualized with VMD (60).

### Molecular Dynamics Simulations (MDS)

To define the interaction between EhSUMO and EhVps32 or EhADH proteins, we performed molecular dynamics simulations (MDS) of EhSUMO and EhVps32 and we took the published data for EhADH (31). The 3D-structure of EhSUMO and EhVPS32 was obtained from their amino acid sequences using ID C4M1C8 and C4M1A5, respectively (uniprot.https://www.uniprotorg), and the I-TASSER server. (https://zhanglab.ccmb.med.umich.edu/I-TASSER). The crystallographic structures that I-TASSER used to obtain the 3D EhSUMO model were: 5GJL solution structure of *Plasmodium falciparum*, IWZ0 solution structure of human SUMO-2 (SMT3B), an ubiquitin-like protein 5XQM structure of SMO1, SUMO homologue of *Caenorhabditis elegans* and 2K8H solution structure of SUMO from *Trypanosoma brucei*. Those used for Ehvps32 model were: 5FD7 and 5FD9 crystal structures of ESCRT-III Snf7 core domain (conformation A and B, respectively) from *Saccharomyces cerevisiae*, 5NNV structure of a *Bacillus subtilis* Sms coiled coil middle fragment, 2GD5 structural basils for budding by ESCRT-III factor ChMP3 from *H. sapiens* and 5NL crystal structure of the two spectrin repeated domains from *E. histolytica*.

MDS were carried out using the NAMD 2.8 software (61). through GPU-CUDA with video cards graphics NVIDIA Tesla C2070/Tesla C2075. The force fields CHARMM22 and CHARMM27 (62) were used to create the topologies for protein and lipids, respectively. The TIP3 model was employed for water molecules. The system was solvated by the *psfgen* software in the VMD program (60). By this software, the NAMD program added water molecules and sodium ions to neutralize the system: one sodium ion was added to EhSUMO with 29329 water molecules; and 14 sodium ions with 8,719 water molecules to neutralize the system for EhVps32. Both systems were minimized for 1,000 steps followed by equilibration, under constant temperature and pressure (NPT) conditions for one ns with protein and lipid atoms restrained. Afterward, 200-ns-long MDS was run, considering EhSUMO and EhVPS32 proteins as soluble, without position restraints under periodic boundary conditions and using an NPT ensemble at 310 K. Particle mesh Ewald technique was calculated for the electrostatic interactions method (63). Nine Å cut-off was used for the van der Waals interactions. The time step was set to 2.0 fs, and the coordinates were saved for analyses every one ps; 200 ns of MDS was carried out for both proteins, then, protein-protein docking calculations were performed using different conformers through 200 ns of MDS. Simulations were performed in the Laboratory of Molecular Modeling and Bioinformatics of the Facultad de Ciencias Químico Biológicas de la Universidad Autónoma de Sinaloa and the Hybrid Cluster Xiuhcoatl (http://clusterhibrido.cinvestav.mx) of CINVESTAV-IPN, México.

The structural analysis from the MDS, the root mean square fluctuation (RMSF), and the radius of gyration (Rg), as well as the snapshots used for docking analysis, were obtained and analyzed with the grcarma software (64). Root mean square deviation (RMSD) was normalized using the SigmaPlot 12.0 software. The protein– protein docking studies were calculated employing different conformers with Cluspro server (65,66). Molecular graphics were performed in SigmaPlot 12.0 and all 3D-structures visualization was performed by VMD (60).

### Cloning of the *E. histolytica SUMO* gene (*EhSUMO)*

To clone *EhSUMO* gene, the full DNA sequence of 345 bp was PCR-amplified, using the following specific primers: sense 5’-CCGGTACCATGTCTAATCAACCACAATATGGAATTAAATC-3’ and antisense 5’-CCGGATCCTTATTTGATGTATTGAAGGTATTGAGTATTAAAAAGA-3’, in a mixture containing 10 mM dNTPs, 100 ng of *E. histolytica* genomic DNA or cDNA as template, and 2.5 U of *Taq* DNA polymerase (Gibco). PCR assay was carried out for 35 cycles (1 min at 94*°*C, 30 sec at 59*°*C, and 40 sec at 72*°*C.) The sense oligonucleotide contains a *KpnI* restriction site, while the antisense oligonucleotide harbors a *BamH1* restriction site. The full-length gene was cloned in pCold II DNA plasmid, which conferred a histidine tag (67). As positive control we use primers to amplify the *EhGATA* gene (68).

### Expression and purification of recombinant protein, and generation of anti-EhSUMO antibodies

*Escherichia coli* BLI21 (pLysS) bacteria were transformed with the pCold/*EhSUMO*, containing the full open reading frame of the *EhSUMO* gene to produce a His-tagged EhSUMO recombinant protein (rEhSUMO). The His-rEhSUMO protein was purified with cobalt beads in an imidazole gradient and used to subcutaneously and intramuscularly inoculate Wistar rats (50 µg emulsified in Titer-Max Classic adjuvant, 1:1) (Sigma), to generate α-EhSUMO polyclonal antibodies. Two more doses (50 µg) were injected at 15-days intervals, and animals were bled to obtain the immune serum. Pre-immune serum was also obtained, before immunization.

### Western blot experiments

Trophozoites lysates (35 µg), or purified rEhSUMO were electrophoresed in 15% sodium dodecyl sulfate polyacrylamide gels (SDS-PAGE), transferred to nitrocellulose membranes and probed with rat α-EhSUMO (1:2,000), mouse α-EhVps32 (1:1,000), rabbit α-EhADH (1:1,000) (34), rabbit α-EhCPADH (1:35,000) (41), mouse α-actin (1:3,000) kindly donated by Dr. José Manuel Hernández (Cell Biology Department, CINVESTAV) and mouse α-EhPCNA (1:500) antibodies (69). Membranes were washed, incubated with α-rat, α-mouse and α-rabbit HRP-labeled secondary antibody (Sigma, 1:10,000), and revealed with ECL Prime detection reagent (GE-Healthcare), according to the manufacturer’s instructions.

### Laser confocal microscopy assays

Trophozoites were grown on coverslips, fixed with 4% paraformaldehyde at 37°C for 1h, permeabilized with 0.5% Triton X-100 and blocked with 10% fetal bovine serum in PBS. Then, cells were incubated at 4°C for overnight (ON) with either α-EhSUMO (1:100) or α-EhVps32 (1:100) antibodies labeled with Alexa-555 or Pacific Blue kit (Molecular Probes-Thermo Fisher), respectively, or with rabbit α-EhADH (1:100) antibodies. After extensive washing, samples were incubated for 30 min at 37°C with α-rabbit FITC-labeled secondary antibody (1:100). Fluorescence was preserved using Vectashield antifade reagent (Vector), examined through a Carl Zeiss LMS 700 confocal microscope, in laser sections of 0.5 μm and processed with ZEN 2009 Light Edition Software (Zeiss). To evaluate the colocation between proteins, fluorescence intensity was quantified from at least 30 confocal images, using the ImageJ 1.45v software and the JACoP plugin.

### Phagocytosis assays

Trophozoites were incubated at 37°C with human erythrocytes (1:25 ratio) for different times (0, 2, 5, 15, 30, and 60 min) and processed for immunofluorescence and western blot assays (70). Three independent experiments for phagocytosis were performed and the number of erythrocytes per trophozoite was obtained from 100 trophozoites.

### Immunoprecipitation assays

Trophozoites were lysed in the presence of 10 mM Tris-HCl, 50-mM NaCl, and proteases inhibitors, by freeze-thawing cycles and vortexing. Immunoprecipitation assays were performed using 200 μl of protein G-agarose (Invitrogen) and α-EhSUMO antibody as described (14). Immunoprecipitates were analyzed by western blot assays using α-EhSUMO, α-EhADH α-EhVps32, and α-actin antibodies, as described above.

### Silencing assay

The knock-down (KD) of *EhSUMO* was performed using the bacterial expression of double-stranded RNA (dsRNA) and parasite soaking experiments as described (39). Briefly, the 345 bp from the 5’-end of the *Ehsum*o gene were PCR-amplified using the following primers: sense 3’-CCGAGCTCATGTCTGACGCACAACATTCA-5’ and antisense 3’-CCGGTACCTTAAAACCCACCAACTTGATTCAT-5’. Then, the amplicon was cloned into pJET1.2/blunt plasmid and subcloned into pL4440 plasmid, using the *Sac1* and *Kpn1* restriction sites. PCR, restriction analysis and DNA sequencing were performed to verify the resulting pL4440-*EhSUMO* plasmid. The competent RNase III-deficient *E. coli* strain HT115 were transformed with the pL4440-*EhSumo*. Bacteria were grown at 37°C in LB broth for plasmid construction or 2YT broth for dsRNA expression, in the presence of ampicillin (100 mg/ml) and tetracycline (10 mg/ml) (71). The expression of *EhSUMO*-dsRNA was induced with 2-mM isopropyl β-D-1-thiogalactopyranoside ON at 37°C. Then, bacterial pellet was mixed with 1 M ammonium acetate and 10 mM EDTA, incubated with phenol:chloroform:isoamyl alcohol (25:24:1) and centrifuged. The supernatant was mixed with isopropanol, centrifuged, and the nucleic acid pellet was washed with 70% ethanol. DNase I (Invitrogen) and RNase A (Ambion) were added to eliminate ssRNA and dsDNA molecules. *EhSUMO*-dsRNA was washed again with isopropanol and ethanol, analyzed by agarose gel electrophoresis and concentration determined by spectrophotometry. Lastly, purified *EhSUMO*-dsRNA (10 μg/ml) molecules were added to trophozoites (3.0×10^4^) in TYI-S-33 complete medium and cultures were incubated at 37°C for 24, 48, 72, 96, and 120 h. At 24 h was the time when silencing of the EhSUMO protein was visualized and analyzed by western blot assays and confocal microscopy. All subsequent experiments were done at this time. Trophozoites growing under standard conditions (without dsRNA), were used as controls.

## Statistical analysis

Statistical analyses were performed by t-Student test, using GraphPad Prism 5.0 software. The scores showing statistically significant differences are indicated with asterisks in the graphs. The corresponding *p* values are indicated in figure legends.

### Ethics statement

The Centre for Research and Advanced Studies (CINVESTAV) fulfil the standard of the Mexican Official Norm (NOM-062-ZOO-1999) “Technical Specifications for the Care and Use of Laboratory Animals” based on the Guide for the Care and Use of Laboratory Animals “The Guide,” 2011, NRC, USA with the Federal Register Number BOO.02.03.02.01.908, awarded by the National Health Service, Food Safety and Quality (SENASICA) belong to the Animal Health Office of the Secretary of Agriculture, Livestock, Rural Development, Fisheries and Food (SAGARPA), an organization that verifies the state compliance of such NOM in Mexico. The Institutional Animal Care and Use Committee (IACUC/ethics committee) from CINVESTAV as the regulatory office for the approval of research protocols, involving the use of laboratory animals and in fulfillment of the Mexican Official Norm, has reviewed and approved all animal experiments (Protocol Number 0505-12, CICUAL 001).

## Acknowledgments

The authors want to express their gratitude to LANCAD for the supercomputer time support.

